# Architectural traits constrain the evolution of unisexual flowers and sexual segregation within inflorescences: an interspecific approach

**DOI:** 10.1101/356147

**Authors:** Rubén Torices, Ana Afonso, Arne A. Anderberg, José M. Gómez, Marcos Méndez

## Abstract

Male and female unisexual flowers have repeatedly evolved from the ancestral bisexual flowers in different lineages of flowering plants. This sex specialization in different flowers often occurs within inflorescences. We hypothesize that inflorescence architecture may impose a constraint on resource availability for late flowers, potentially leading to different optima in floral sex allocation and unisexuality. Under this hypothesis we expect that inflorescence traits increasing the difference in resource availability between early and late flowers would be phylogenetically correlated with a higher level of sexual specialization. To test this hypothesis, we performed a comparative analysis of inflorescence traits (inflorescence size, number of flowers and flower density) in the sunflower family, which displays an extraordinary variation in floral sexual specialization at the inflorescence level, i.e. hermaphroditic, gynomonoecious and monoecious species. We found that species with a complete sex separation in unisexual flowers (monoecy) had significantly denser inflorescences. Furthermore, those species arranging their flowers in denser inflorescences also showed greater differences in the size of early and late fruits, a proxy of resource variation between flowers. Our findings support the idea that floral sexual specialization and consequently sexual segregation may be the consequence of different floral sex allocation optima driven by the sequential development of flowers that results in a persistent resource decline from earlier to later flowers.

## Introduction

Most angiosperms are hermaphrodite, i.e., they produce bisexual or 'perfect' flowers bearing both functional pollen grains and ovules. Combining both sexual functions in the same flower reduces direct reproduction costs, such as sharing attractive structures for pollinators [1], and acts as an insurance against vagaries in the mating environment by allowing selfing [2]. Thus, evolutionary biologists have wondered, since Darwin [3], what favours non-hermaphroditic sexual systems, where flowers of different sex -male, female, or bisexual-are combined in the same or different individuals of a population. Darwin [3] interpreted the evolution of unisexual flowers as a means of reducing selfing. Nevertheless, the simultaneous presence of self-incompatibility and unisexual flowers in many angiosperms [4] suggests that avoiding selfing is not the only driver of sexual specialization in flowers. Alternative explanations for the evolution of sexual specialization are the avoidance of interference between sexual functions [5,6] and optimal resource allocation [6–8].

Any evolutionary explanation of sexual specialization of flowers should consider that sexes are often segregated within inflorescences due to two constraints set by inflorescence architecture. The first constraint is ontogenetic [9–11]: because flowers within inflorescences usually develop sequentially, early flowers usually flower and ripen fruits first and pre-empt resources for late flowers. The second constraint is strictly positional [11] and results in a persistent limitation of resources at certain flower positions inherent to the architecture of inflorescence axes. Both constraints provide a general proximate mechanism for sexual selection, because they can strongly influence the mating environment or resource availability experienced by individual flowers [11–13]. As resource availability may differentially affect male and female performance [14,15], this resource gradient across flowers within an inflorescence could also lead to differential sexual specialization in female or male functions at different floral positions.

On theoretical grounds, a female-biased floral allocation can be predicted in those floral positions in the inflorescence with higher resource availability, whereas a male-biased allocation will be expected in positions with less resources [12]. Intraspecific observational [11] and experimental evidence [16,17] supports this idea, at least for plants with elongated inflorescences. For instance, *Solanum hirtum* produces inflorescences with bisexual flowers, but the late (distal) ones are labile, becoming male in resource-depleted plants [17]. Nevertheless, it remains poorly understood whether these intraspecific plastic responses in sexual specialization and segregation within inflorescences can actually become fixed during the evolution of a lineage [11,18], giving rise to the sexual segregation within inflorescences observed in many angiosperm families. The exception is a pioneer comparative study of Solanum that found how plastic responses in the production of male unisexual flowers are ancestral to fixed position effects [19]. Here, we take this comparative approach a step further, by using Asteraceae to test at a family level whether the degree of sexual specialization is phylogenetically correlated with inflorescence architectural traits which may lead to a resource decline from early to late flowers.

We hypothesize that three architectural traits of inflorescences, namely larger size, and higher flower number and density increase resource competition among flowers and likely constrain resource availability for late flowers, potentially leading to male-biased floral sex allocation and unisexuality. Testing this hypothesis at an interspecific level faces two challenges. First, inflorescences may differ impressively across species in size, number of flowers, fruit size and flower size, even within the same main inflorescence type [20–23]. Second, these traits do not necessarily covary across species, because inflorescence size and flower number are under selection by several ecological drivers, such as pollinators [24–28], seed predators [29], or altitude and geographic ranges [30]. Consequently, the diversity of trait combinations across species could obscure the detection of relationships between inflorescence traits and floral sexual specialization. In particular, a negative covariation between flower size and number [31–33] could lead to similar levels of resource competition among flowers/fruits in inflorescences having different number of flowers, since an increase in the number of flowers is commonly associated with a decrease in flower and fruit size. Thus, flower density, in which flower size and inflorescence size are taken into account, may be a more reliable indicator of the strength of resource competition among flowers within an inflorescence than the number of flowers per inflorescence or inflorescence size alone.

In this study, we compared species of the sunflower family showing different levels of floral sexual specialization to test whether they show different inflorescence traits. We then assessed how these inflorescence traits correlate with resource differences between flowers within the same inflorescence. The sunflower family (Asteraceae) is a suitable model for testing this hypothesis. First, all Asteraceae share the same basic inflorescence architecture, the head or capitulum [34], which mainly follows a centripetal pattern in floral development and blooming [35,36]. Second, three different sexual systems are common in Asteraceae: hermaphroditism, gynomonoecy and monoecy (Fig. 1). From the ancestral hermaphroditism, monoecy likely evolved through a gynomonoecious intermediate [37]. Third, a clear pattern of sexual specialization is present within inflorescences in non-hermaphroditic species. Namely, gynomonoecious species produce female flowers at the outermost positions and bisexual ones in the inner positions, while monoecious species show female flowers in the outermost positions and male flowers in inner positions (Fig. 1). Finally, there is anatomical [38], physiological [39] and experimental evidence that architectural constraints occur within heads, and both outer flowers and positional effects can limit the available resources to late-blooming, inner flowers [18].

**Figure 1.**
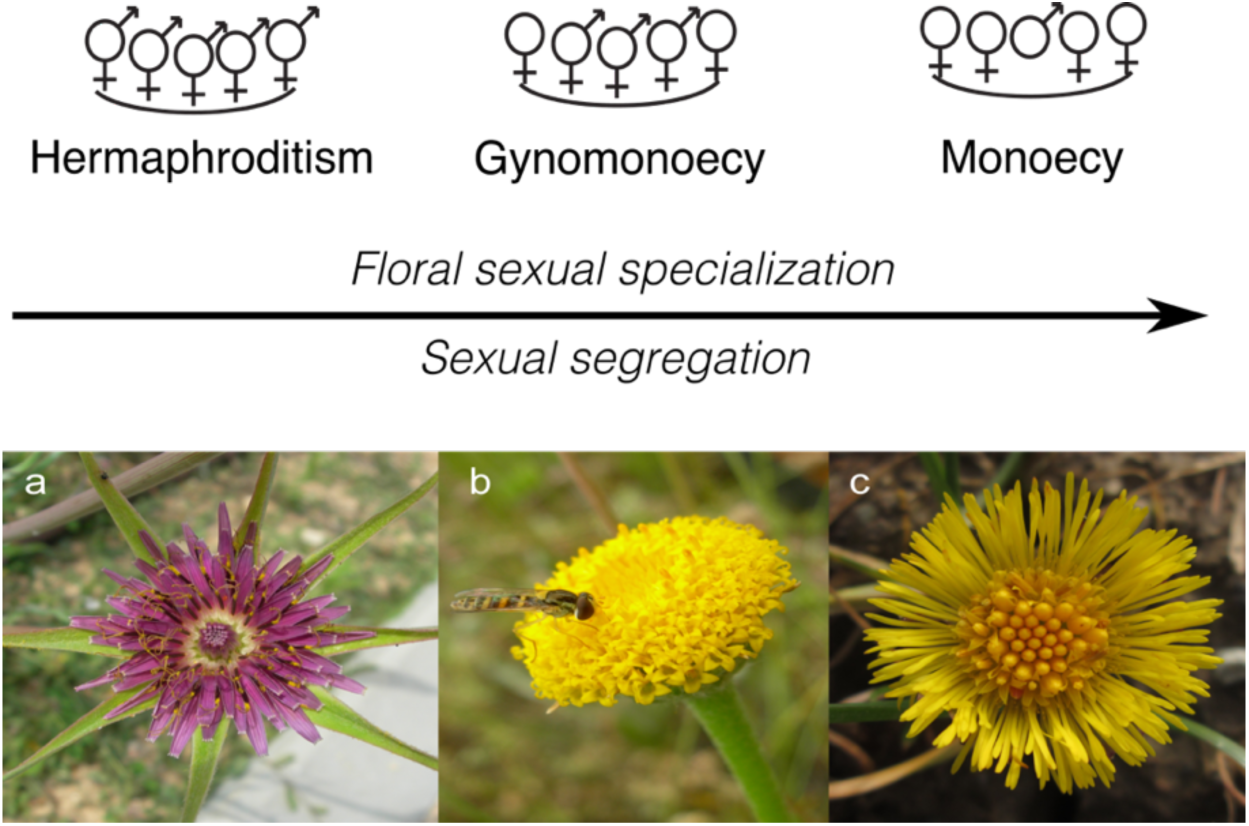
Sexual specialization in the Asteraceae inflorescences. Three different types of functionally hermaphroditic inflorescences and individuals can be observed in the Asteraceae: hermaphroditic, gynomonoecious and monoecious. Allocation of gametes to different flowers shows a gradient in floral sexual specialization from hermaphroditic species (only bisexual flowers) to gynomonecious species (female and bisexual flowers) and to monoecious species (female and male flowers). In Asteraceae, sexual specialization (i.e., bi- vs. unisexuality in flowers) and sexual segregation within inflorescences occur in concert. Lower panels show representative species from **(a)** hermaphroditic heads (*Tragopogon porrifolius* L.), **(b)** gynomonoecious heads (*Anacyclus valentinus* L.), and **(c)** monoecious heads (*Tussilago farfara* L.).

In this paper, we followed a comparative approach to study the role of architectural constraints in the evolution of floral sexual specialization and sexual segregation within inflorescences in the Asteraceae. Specifically, we assessed how inflorescence traits (namely, inflorescence size, number of flowers, and flower density) were associated with different levels of floral sexual specialization and sexual segregation represented by hermaphroditic, gynomonoecious and monoecious species (Fig. 1). In addition, we assessed how these inflorescence traits affected variation in fruit size within the inflorescences, as a proxy for the resource gradient between flower positions [9,40]. Finally, we tested whether female flowers in an inflorescence produce larger fruits than bisexual ones, as would be expected if sexual specialization 'releases' female flowers from expending resources on male structures and compensated the lack of the male sex function by producing larger fruits than bisexual flowers.

## Methods

### Study species

The Asteraceae is the largest family of angiosperms, with over 1,500 genera and 25,000 species, and a worldwide distribution [34]. All Asteraceae share the same basic inflorescence, the head or capitulum, a dense indeterminate inflorescence where all the flowers are sessile and attached to a common receptacle (Fig. 1). Heads represent the basic pollinator attraction unit [20,21]. Different degrees of sexual segregation within the heads can be observed among species of the family (Fig. 1). From hermaphroditism to monoecy, a floral sexual specialization occurring in individual flowers. The evolutionary transition between them occurs through a gynomonoecious intermediate that bears both female and bisexual flowers [37].

We included a total of 91 species in our study, including 42 hermaphroditic, 28 gynomonoecious and 18 monoecious species (Supplementary Table S1). The species were studied on material from two herbaria: the Asteraceae collection of the Swedish Natural History Museum Herbarium (S), and the Herbarium of the University of Coimbra (COI). Herbarium sampling allowed a comprehensive sampling of Asteraceae diversity, since different evolutionary lineages coming from different continents and biomes were sampled. We sampled specimens from 26 tribes and 7 subfamilies (including the “Heliantheae alliance”). Hermaphroditism is the ancestral condition in the family and also in most of the early diverging lineages (Supplementary Fig. S1; [37]). Once gynomonoecy evolved at the origin of the subfamily Asteroideae, it remained as the ancestral condition for most of the tribes of this large clade (Fig. S1). This evolutionary history of sexual systems was taken into account when selecting the species to be sampled and we put a strong emphasis in sampling representatives of all sexual systems at the early and late diverging lineages, as well as in sampling basal and derived hermaphroditic and gynomonoecious species. All monoecious species were almost always derived from gynomonoecious lineages in the subfamily Asteroideae where all monoecious species evolved in this family with very rare exceptions.

### Inflorescence and fruit traits

Three traits were measured at an inflorescence level in the 91 species: (i) inflorescence size, measured as the head diameter in mm; (ii) the number of flowers per inflorescence; and (iii) flower density, calculated as the ratio between the number of flowers and the area of each head. Flower density was used as a measure of floral aggregation and may provide a better proxy of resource competition between flowers within the head than head size or number of flowers per head.

To minimize specimen damage, we sampled only those specimens with mature fruits, i.e. fruiting heads, in which we measured inflorescence and fruit traits. Fruiting heads in Asteraceae usually retain the size and structure of the inflorescence and therefore they can be used to describe inflorescence traits such as size and number of flowers. Otherwise, we assume that any change in size during the ripening of the fruiting head is proportional to head size. First, we searched those specimens belonging to the species included in the phylogenetic supertrees published for this family [34,41]. Second, we selected herbarium specimens with enough mature fruiting heads and maintaining their original positions within heads. For each species, one specimen having fruiting heads at desirable conditions was selected, and at least one capitulum was sampled. We sampled the specimen in which fruiting head removal caused the least damage possible to the specimen. Following this procedure, 109 herbarium specimens were initially sampled, and from 91 of them we could collect inflorescence traits (Supplementary Table S1).

All fruiting heads were dissected to separate all fruits in their relative positions within heads from the outermost to the innermost positions. When necessary, heads were placed in water with a detergent for rehydration and to reduce damage to the head. Data were collected for 70 species. Heads and fruits were measured using pictures taken with a tripod-stabilized digital camera. Fruit area of more than 2,700 fruits were measured as using Image J 1.54s software [42]. Although low intraspecific sample sizes may lead to increased type I error in comparative studies, this effect is important only when coupled with high intraspecific variation [43]. However, when the range of taxa studied is wide, variation across species is usually much greater than variation within species. In our study, fruit size varied between species investigated, the largest fruits were more than 100-fold larger than the smallest fruits, whereas no single species showed such a degree of variation in fruit size.

### Statistical analyses

#### Relationship between inflorescence traits and floral sexual specialization

We used phylogenetic generalized least squared (PGLS) models [44,45] to explore the relationship between the degree of floral sexual specialization (hermaphroditism, gynomonoecy or monoecy) and inflorescence traits. All models were evaluated under both an adaptive model (OU, Ornstein-Uhlenbeck model) and a neutral model of evolution (BM, Brownian motion model) [46]. PGLS were fitted using the “ape” [47] and “geiger” packages [48] in R. The fittest model for each combination of variables was selected using a likelihood ratio test comparing BM and OU models. For all fitted models, the OU model had a higher goodness of fit than the BM (results not shown). Therefore, we present only the results fitted under an OU model. The phylogenetic relationship between the species included in the analyses was considered by using the phylogenetic supertree published for the Asteraceae [41], adding a calibration to include branch lengths [49]. The root of this tree was previously scaled to 1 for all the analyses. Specific comparisons between hermaphroditism, gynomonoecy and monoecy were explored using the marginal means, using the ‘lsmeans’ package [50] in R [51], which can be defined as a linear combination of the estimated effects from a linear model.

#### Relationships between fruit size, inflorescence traits and floral sexual specialization

We explored how fruit size variation within inflorescences, as a proxy of the resource gradient between flower positions, was related to inflorescence traits and floral sexual specialization, using the meta-analytical effect size to get a standardized measure of the magnitude of the difference among the size of the outer and inner fruits (fruit size difference, hereafter FSD). A random-effects meta-analysis was used. Effect sizes were calculated using the ‘meta’ package in R [52].

The correlation between FSD and fruit size with the degree of floral sexual specialization was explored using PGLS models. FSD and fruit size were the response variables, whereas the degree of floral sexual specialization was the predictor variable. All models were evaluated under both an OU and a BM model (see above).

In addition, we assessed the allometric relationships of inflorescence traits (head diameter, number of flowers and flower density) with FSD and outer and inner fruit sizes. These allometric relationships were tested by fitting PGLS models. FSD and fruit size were the response variables, and the three inflorescence traits were the predictor variables. For those significant correlations, we estimated phylogenetic reduced major axis regressions and tested whether slopes were significantly different from one using the ‘phyl.RMA’ function included in the ‘phytools’ package [53]. All variables were log-transformed before analysis.

#### Relationship between floral sexual specialization and fruit size

We explored whether unisexual flowers produced larger fruits than bisexual flowers. We assessed the effect of floral sexual specialization (female vs. bisexual flowers) using only the outer fruits because strictly female flowers are found only in these positions. In addition, given the strong effect of inflorescence traits on fruit size (see Results), we included in the model the density of flowers, to control for the size of the inflorescence and the number of flowers. We fitted two PGLS models, where fruit size was the response variable and flower sex was included as a predictor categorical variable. Flower density was included as a continuous predictor variable. The only difference between both models was the inclusion of an interaction term between the sex of the flower and the flower density. The model including an interaction did not perform better and was thus dropped. Fruit size and flower density were log-transformed. We evaluated this model under OU and BM correlation structures.

## Results

### Inflorescence traits and floral sexual specialization

Inflorescence traits differed significantly among the degrees of floral sexual specialization, i.e., hermaphroditism, gynomonoecy, and monoecy (Table 1). Hermaphroditic and gynomonoecious species displayed larger inflorescences than monoecious species (Fig. 2a). Gynomonoecious species had significantly more flowers per inflorescence than hermaphroditic and monoecious species (Fig. 2b). Nevertheless, flower density was correlated with the degree of floral sexual specialization, increasing from hermaphroditic through gynomonoecious to monoecious species (Fig. 2c).

**Table 1.**
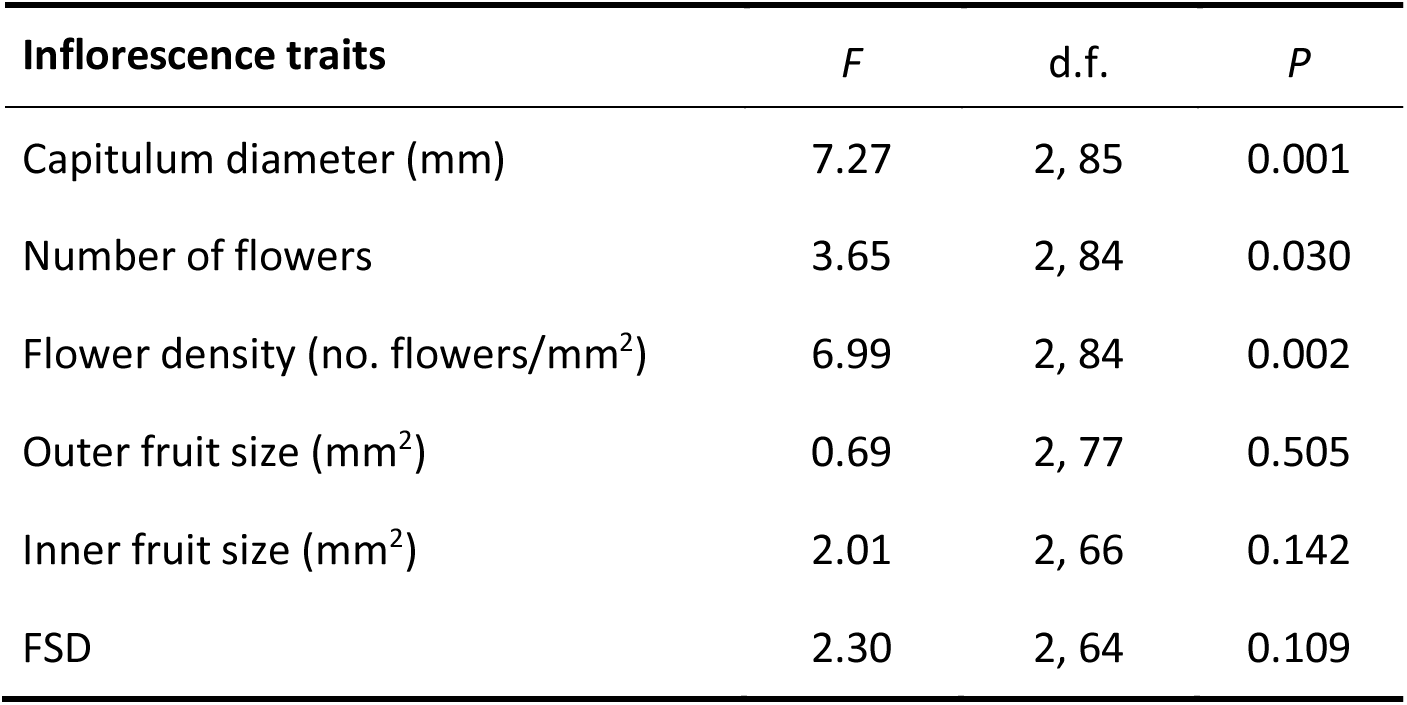
Differences in inflorescence and fruit traits between inflorescences with different degree of floral sexual specialization. *F* and *P* values were obtained after a deviance analysis of the phylogenetic generalized linear models fitted for each inflorescence trait, with the degree of floral sexual specialization within inflorescence (hermaphroditism, gynomonoecy or monoecy) as the main factor. FSD is the standardized fruit size difference between outer and inner fruits measured as the meta-analytical effect size.

**Figure 2.**
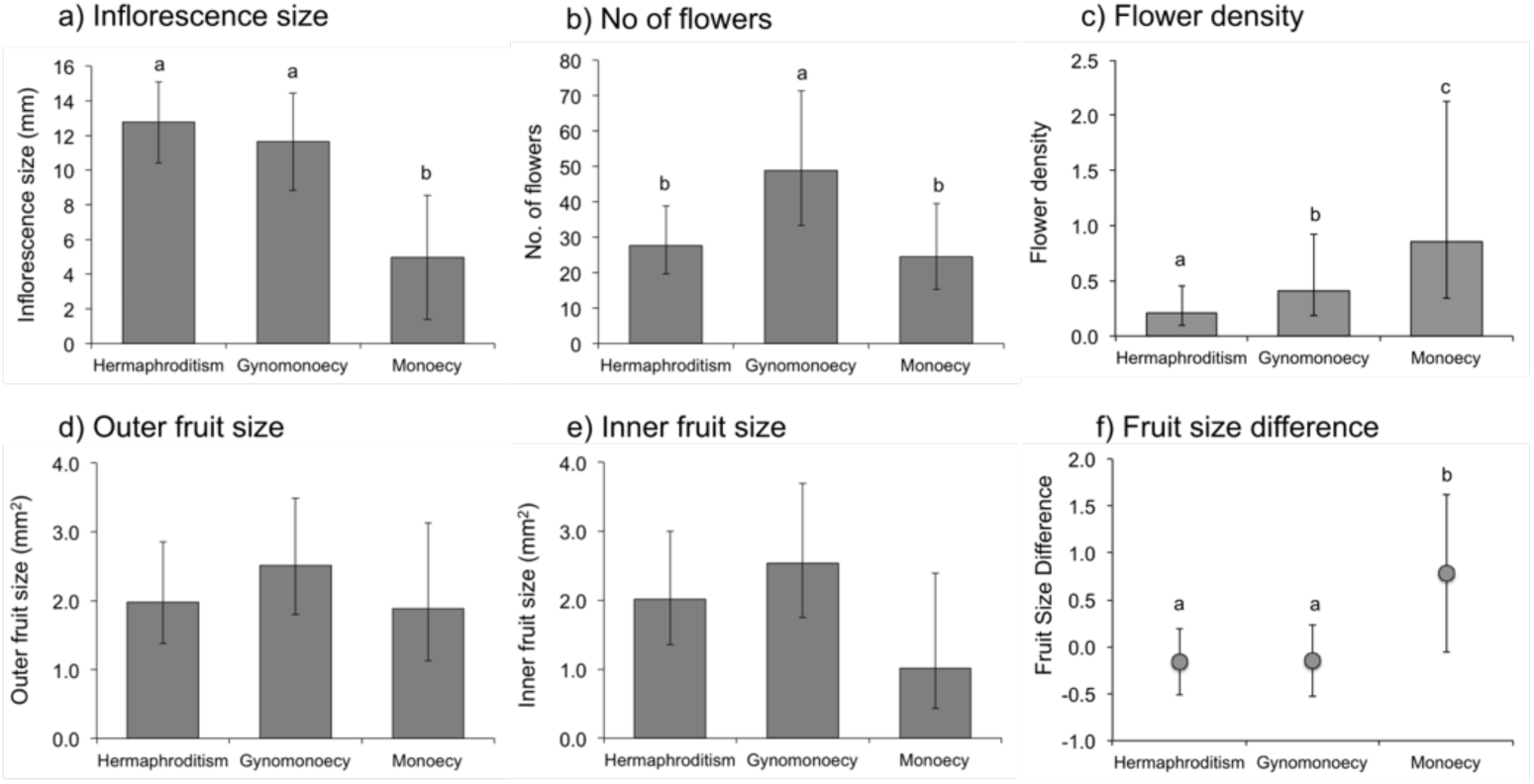
Inflorescence and fruit traits of different sexual systems. Phylogenetically controlled least-squares means (± 95% confidence interval) of **(a)** inflorescence size (mm), **(b)** number of flowers, **(c)** flower density (no. flowers / mm^2^), **(d)** outer fruit size, **(e)** inner fruit size, and **(f)** Fruit Size Difference (FSD) for different levels of sexual systems representing increasing levels of floral sexual specialization within inflorescences: hermaphroditism, gynomonoecy and monoecy. FSD is the standardized fruit size difference between outer and inner fruits measured as the meta-analytical effect size. Means sharing the same superscript letter were not significantly different at the *P* < 0.05 level.

Inflorescence size was significantly correlated to the other two inflorescence traits (Supplementary Table S2). The number of flowers increased disproportionally with an increase in inflorescence size, whereas flower density disproportionally decreased with an increase in inflorescence size, measured as head diameter (Supplementary Table S3). Number of flowers and flower density were only marginally correlated (Supplementary Table S2), and flower density increased proportionally with an increased number of flowers (Supplementary Table S3).

### Fruit size variation within inflorescences

Fruit size was not statistically different between hermaphroditic, gynomonoecious and monoecious species, either for outer or inner positions (Table 1; Fig. 2d,e). Two different relationships between fruit size and inflorescence traits were observed, independently of position in the head. First, fruit size significantly increased with an increase in inflorescence size (Fig. 3a). The phylogenetic RMA slopes were significantly higher than one (outer fruits: *b* = 1.34, t = 3.16, d.f. = 68.9, *P* = 0.002; inner fruits: *b* = 1.40, t = 3.64, d.f. = 57.4, *P* < 0.001) indicating a disproportionate increase in fruit size with an increase in inflorescence diameter (Fig. 3a). Second, fruit size decreased with an increase in flower density (Fig. 3c). The phylogenetic RMA slopes were significantly < 1.0 (outer fruits: *b* = −0.79, t = 3.31, d.f. = 61.4, *P* = 0.002; inner fruits: *b* = −0.82, t = 2.78, d.f. = 51.7, *P* = 0.008). Therefore, fruit size decreased at a lower rate than the increase in flower density (Fig. 3c). Fruit size was not statistically correlated with flower number (Fig. 3b).

**Figure 3.**
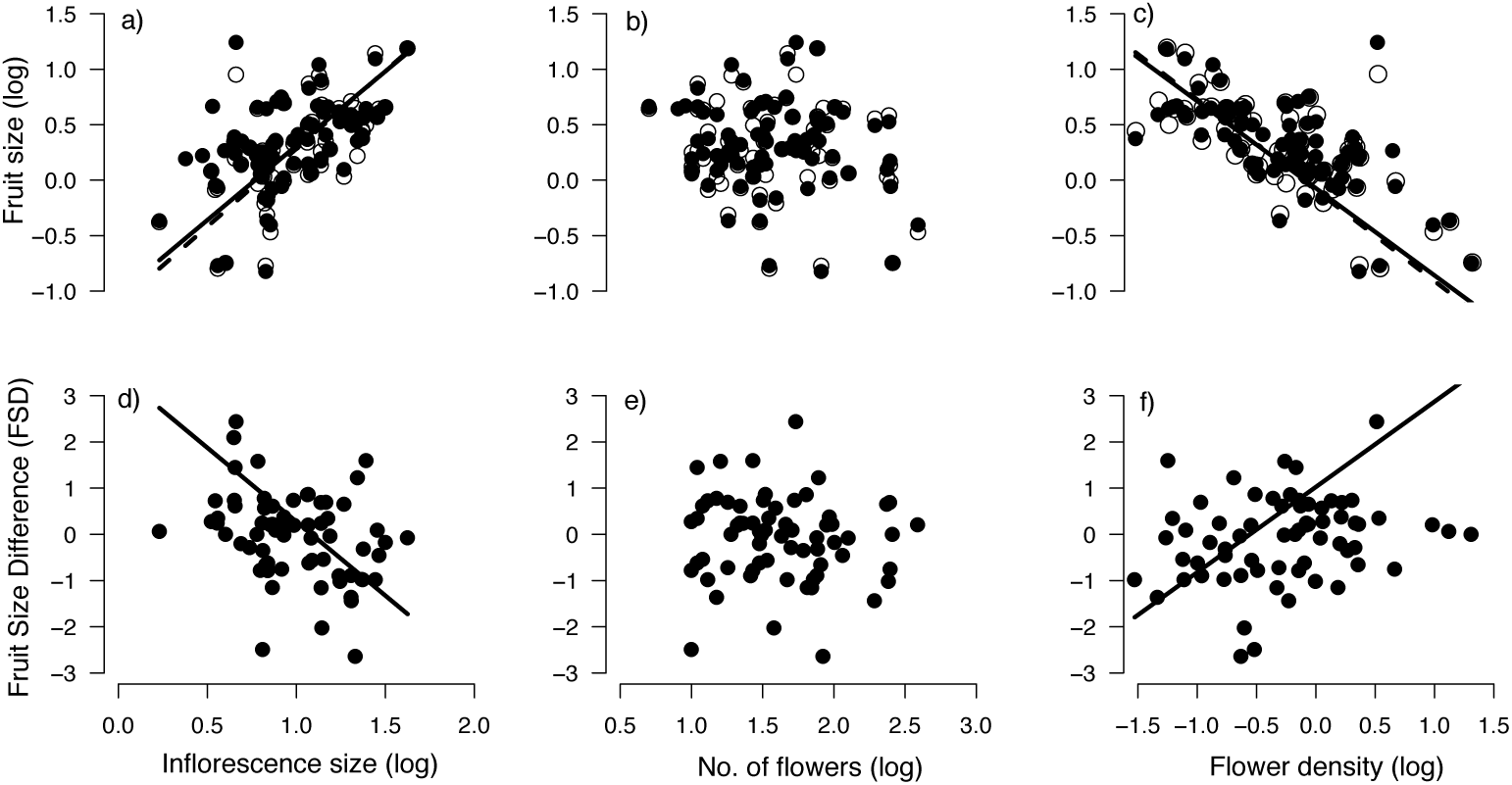
Allometric relationships between fruit size and inflorescence traits. Phylogenetic RMA regressions between fruit size and **(a)** inflorescence size (mm), **(b)** number of flowers, and **(c)** flower density (no. flowers / mm^2^); and between FSD and **(d)** inflorescence size (mm), **(e)** number of flowers, and **(f)** flower density (no. flowers / mm^2^). In the upper row black dots and solid lines represent outer fruits whereas white dots and dashed lines indicate inner fruits. Lines represent RMA slopes. Phylogenetic standard regression slopes are showed in Supplementary Table 4.

FSD, the standardized fruit size difference, significantly decreased with an increase in inflorescence size (Fig. 3d; *b* = −3.20, t = 10.24, d.f. = 62.2, *P* <0.0001), implying that smaller heads showed a higher difference between outer and inner fruits. FSD did not show any significant relationship with the number of flowers (Fig. 3e), although it significantly increased with flower density (Fig. 3f), revealing that an increase in flower density was associated with a higher difference in size between outer and inner fruits (*b* = 1.85, t = 5.28, d.f. = 61.9, *P* <0.0001). The degree of sexual specialization did not significantly affect FSD (Table 1). However, the post-hoc comparison between degrees of sexual specialization showed that monoecious species had significantly larger FSD than hermaphroditic and gynomonoecious species (Fig. 2e). This difference was mainly mediated by the differences among sexual systems in flower density, because the inclusion of flower density as a covariate removed any statistical difference between monoecious species and the other two categories considered (Supplementary Table S5).

### Floral sexual specialization and fruit size

Fruit size decreased with increasing flower density (*F*_1,76_ = 109.03, *P* < 0.0001; Fig. 4). Nevertheless, female flowers produced significantly larger fruits than bisexual flowers after controlling by flower density (*F*_1,76_ = 5.87, *P* = 0.018, n = 78; Fig. 4).

**Figure 4.**
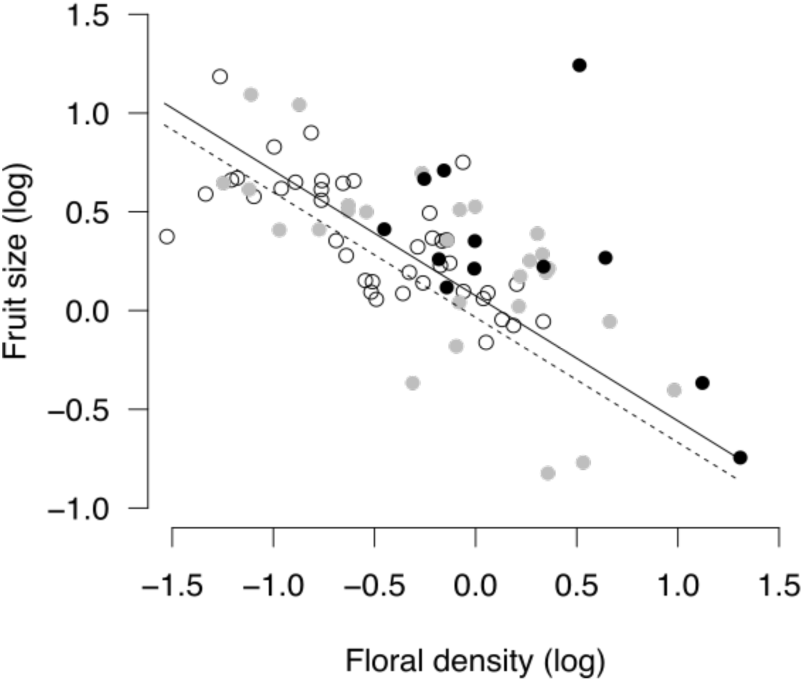
Relationship between fruit size produced by female (filled dots and solid line) and bisexual flowers (white dots and dashed line) and flower density. Female flowers produced by monoecious species are shown as black dots whereas those produced by gynomonoecious species are indicated by grey dots.

## Discussion

Our comparative study across three sexual systems in Asteraceae provided clear support for a phylogenetic association between inflorescence traits, fruit size variation within inflorescences, and the degrees of sexual specialization and segregation. In particular, we observed that monoecious species bore smaller and denser inflorescences than hermaphroditic ones, also showing the largest fruit size difference (FSD) between outer and inner fruits. These results, together with the lack of a correlation between the number of flowers and the degree of sexual specialization, support the idea that floral sexual specialization and consequently sexual segregation within inflorescences of Asteraceae might in part be the result of different sex allocation optima brought about by architecturally mediated persistent resource limitation of the inner flowers.

Although selfing and geitonogamy avoidance has been considered an important factor in the evolution of floral sex specialization and sexual segregation within inflorescences [54], on its own it is unlikely to explain the pattern observed in Asteraceae. The negative effects of geitonogamy are expected to be higher in large inflorescences, where a higher number of flowers can lead to longer floral bouts by pollinators [55]. Thus, many-flowered inflorescences would be expected to show a higher probability of exhibiting unisexual flowers than few-flowered inflorescences. However, in our data set the number of flowers per inflorescence was not clearly related to sexual specialization (Fig. 3b). Moreover, outer seeds should be more outcrossed than inner ones, but evidence is scarce and inconclusive: two species have shown higher outcrossing rate in outer flowers than inner ones [56,57], whereas two other studies have found no differences [58,59]. Finally, the species of this family usually show other mechanisms to avoid selfing such as self-incompatibility [60] and dichogamy (both at the flower and inflorescence levels; [21]). Therefore, whether limiting geitonogamy plays a key role in the evolution of sexual segregation in Asteraceae requires further examination, because empirical evidence remains contradictory.

An alternative to the selfing avoidance hypothesis is that evolutionary transitions between sexual systems might have evolved as a way to optimize gamete packaging. Inflorescence traits, such as the inflorescence size, inflorescence number, but also the number of flowers per inflorescence, are key components of gamete packaging strategies [61,62]. Our results suggest that shifts in the so called ‘inflorescence design’, can have effects at the flower level too. For instance, the increase in flower number is usually negatively correlated with flower size [31–33]. In the Asteraceae, these shifts can have triggered shifts in floral sex expression leading to transitions between hermaphroditism, gynomonoecy and monoecy, whereas the pollination unit keeps their mating opportunities through the retention of both male and female functions. Two ‘inflorescence design’ syndromes has been proposed related to sexual systems for the tribe Inuleae [63] which are also supported by our results. In the first group, including hermaphroditic and gynomonoecious species, attraction units were mostly large solitary or corymbose heads with rather few heads together. In contrast, the attraction unit of the second group, composed of monoecious species only, were clusters of several small heads. Moreover, we observed that although larger inflorescences produced both more flowers and larger fruits in species in our sample, fruit size decreased with flower density. This indicates that resource competition can be the mechanism underlaying the interspecific negative relationships between the number of flowers and fruit size.

Importantly, our study provides insights into how intraspecific size-number trade-offs can translate into negative covariation between traits across species. The consequences of this negative correlation between size and number at the inflorescence level were not the same for all flower positions. We observed that increased flower density led to decreases fruit size, especially at the innermost positions, resulting in a higher FSD (fruit size difference between outer and inner fruits). Flower density might thus amplify the effects of architectural constraints, which pervasively limit resources at the innermost positions. Under these circumstances, theory predicts that a high resource difference between flower positions can cause plants to allocate more resources to their female function in flowers with more resources, and to their male function in resource-depleted flowers [12]. This expectation agrees with the positional pattern observed for the sex of flowers in Asteraceae inflorescences (Fig. 1), where female unisexual flowers consistently appear at those earlier (or outer) floral positions that generally have a higher resource supply, whereas male unisexual flowers are displayed at the later (or inner) positions, which usually are the most resource-limited positions [18,38,39]. Therefore, shifts in inflorescence traits modifying the density of flowers might secondarily cascade to other important floral traits such as flower and fruit size. Specially, flower density might have a role on the evolution of floral sex functions, given its effects on fruit size and thus on the floral female performance.

Under the resource optimization hypothesis, floral sex specialization is expected to entail an improvement in fitness compared to bisexual flowers. Previous studies indicate that dioecious species have larger fruit set than cosexual species [64] and larger seed size [65]. While there is not a formal test on dioecious species of the Asteraceae or in other groups where flowers are no longer the functional unit, our results support that when the inflorescence is the main pollination unit, architectural traits such as flower density may obscure direct comparisons between dioecious and non-dioecious species, given its effect on fruit size. In the Asteraceae, female unisexual flowers from both gynomonoecious and monoecious species had significantly larger fruits than bisexual species, when the confounding effect of flower density was factored out. Thus, our study provides support for an intrinsic advantage of flower specialization at least in terms of female fitness.

Expectations of a negative covariation of traits across species assumes that everything else remains equal. Vamosi *et al*. [65] did not find any difference in seed size between hermaphroditic and monoecious species. However it is unclear if hermaphroditic and monoecious species in their dataset differed in inflorescence traits, such as flower density, which could confound the effect of the floral sexual specialization on fruit and seed size. Positive correlations between inflorescence parts might be found if species differ in resource budget (the big house-big car effect, *sensu* [66]). In addition, additional selective factors on dispersal performance, which usually occur at the fruit level, could indirectly drive shifts on inflorescence design. Our study only revealed the evolutionary drivers in the specialization of floral sex functions after taking into account other inflorescence traits that might otherwise have masked those drivers. This underlies the strength of a comparative approach for understanding the mechanistic basis of the evolution of non-hermaphroditic sexual systems, in the line of Diggle & Miller [19], even when considering phylogenetic scales above the genus level.

### Conclusion

Our results highlight the importance of considering architectural traits to understand phenotypic diversity in modular organisms, which can have important functional consequences ([67] for a review). Architectural constraints may have profound consequences in modular organisms such as plants, influencing how male and female functions perform at different positions within one individual. In particular, the sequential development of inflorescences and asymmetric competition between early and late flowers lead to a gradient in the resource availability experienced by individual flowers within the inflorescences. Thus, a separation of male and female functions in different flowers might evolve not only to maximize mating patterns, but also to optimize resource allocation.

This combination of architectural constraints and selection for optimal sex allocation at the flower level could explain sexual specialization and segregation in other clades beside Asteraceae. Differential access to resources and/or architectural constraints derive from basic anatomical properties of inflorescences and therefore should be general to most plant clades. In fact, in a sample of 127 families showing andro-, gyno-or monoecy, two thirds showed sexual segregation along inflorescences (M. Méndez, unpubl. data), suggesting that inflorescence architecture is a general constraint in the evolution of these sexual systems. Particular aspects of gamete packaging could differ among clades or inflorescence architectures. For example, floral density can have a minor role in elongated inflorescences compared to inflorescence size and flower number. This is why a comparative approach to sexual specialization and segregation has to be aware that all else will rarely remain equal and consider that different solutions may exist to a common problem. Comparative studies including inflorescence traits, sexual expression and fruit size variation across flowers offer great promise in elucidating the generality of this mechanism in the evolution of unisexual flowers in Angiosperms.

## Data accessibility

Data are available at ZENODO: https://doi.org/10.5281/zenodo.2565327.

## Supplementary material

Supplementary tables S1-5 and figure S1 are available online at bioRxiv: https://doi.org/10.1101/356147.

## Acknowledgements

We are especially grateful to John Pannell for the review of a previous version of the manuscript, and to L. DeSoto, S. Castro and F. Sales for their help in different stages of the study. RT was partially supported by the Spanish Ministry of Education (BVA 2010-0375). This preprint has been reviewed and recommended by Peer Community In Evolutionary Biology (https://dx.doi.org/10.24072/pci.evolbiol.100069). We thank the Recommender, Juan Arroyo, and three anonymous reviewers for very helpful comments on previous versions of this paper.

## References

[1] Charnov, E. L., J. Maynard Smith, J. J. Bull. Why be an hermaphrodite? Nature 263 (1976), 125–126.

[2] Wilson, W. G., and L. D. Harder. Reproductive uncertainty and the relative competitiveness of simultaneous hermaphroditism versus dioecy. Am. Nat. 162(2003), 220–241.

[3] Darwin, C. The different forms of flowers on plants of the same species. John Murray, London, UK, 1877.

[4] Bertin, R. I. Incidence of monoecy and dichogamy in relation to self-fertilization in angiosperms. Amer. J. Bot. 80 (1993), 557–560.

[5] Charnov, E. L. Simultaneous hermaphroditism and sexual selection. Proc. Natl. Acad. Sci. USA 76 (1979), 2480–2484.

[6] Willson, M. F. Sexual selection in plants. Am. Nat. 113 (1979), 777–790.

[7] Janzen, D. H. A note on optimal mate selection by plants. Am. Nat. 111, (1977) 365–371.

[8] Lloyd, D. G. Selection of combined versus separate sexes in seed plants. Am. Nat. 120 (1982), 571–585.

[9] Lee, T. D. Patterns of fruit and seed production. Pp 179–202 in J. Lovett-Doust and L. Lovett-Doust. Plant reproductive ecology: patterns and strategies. Oxford University Press, New York, 1988.

[10] Diggle, P. K. Architectural effects and the interpretation of patterns of fruit and seed development. Annu. Rev. Ecol. Syst. 26 (1995), 531–552.

[11] Diggle, P. K. Architectural effects on floral form and function: a review. Pp. 63–80 in T. Stuessy, E. Hörandl, and V. Mayer, eds. Deep morphology: toward a renaissance of morphology in plant systematics. Ganter Verlag, Königstein, Germany, 2003.

[12] Brunet, J., and D. Charlesworth. Floral sex allocation in sequentially blooming plants. Evolution 49 (1995), 70–79.

[13] Harder, L. D., and P. Prusinkiewicz. The interplay between inflorescence development and function as the crucible of architectural diversity. Ann. Bot. 112 (2013), 1477–1493.

[14] Freeman, D. C., K. T. Harper, and El. L. Charnov. Sex change in plants: old and new observations and new hypotheses. Oecologia 47 (1980), 222–232.

[15] Lau, T.-C., and A. G. Stephenson. Effects of soil nitrogen on pollen production, pollen grain size, and pollen performance in Cucurbita pepo (Cucurbitaceae). Amer. J. Bot. 80 (1993), 763–768.

[16] Emms, S. K. Andromonoecy in Zigadenus paniculatus (Liliaceae): Spatial and temporal patterns of sex allocation. Am. J. Bot. 80 (1993), 914–923.

[17] Diggle, P. K. The expression of andromonoecy in Solanum hirtum (Solanaceae): phenotypic plasticity and ontogenetic contingency. Am. J. Bot. 81 (1994), 1354–1365.

[18] Torices, R., and M. Méndez. Fruit size decline from the margin to the center of capitula is the result of resource competition and architectural constraints. Oecologia 164 (2010), 949–58.

[19] Diggle, P. K., and J. S. Miller. Developmental plasticity, genetic assimilation, and the evolutionary diversification of sexual expression in Solanum. Am. J. Bot. 100 (2013), 1050–1056.

[20] Leppik, E. E. The evolution of capitulum types of the Compositae in the light of insect-flower interaction. Pp 61–89 in V. H. Heywood, J. B. Harborne, B. L. Turner, eds. The biology and chemistry of the Compositae. Vol. 1. Academic Press, London, UK, 1977.

[21] Burtt, B. L. Aspects of diversification in the capitulum. Pp. 41–59 in V. Heywood, J. B. Harborne, B. L. Turner, eds. The biology and chemistry of the Compositae. Vol. 1. Academic Press, London, 1977.

[22] Wyatt, R. Inflorescence architecture: how flower number, arrangement, and phenology affect pollination and fruit-set. Am. J. Bot. 69 (1982), 585–594.

[23] Harder, L. D., C. Y. Jordan, W. E. Gross, and M. B. Routley. Beyond floricentrism: the pollination function of inflorescences. Plant Species Biol. 19 (2004), 137–148.

[24] Willson, M. F., and B. J. Rathcke. Adaptive design of the floral display in Asclepias syriaca L. Am. Midl. Nat. 92 (1974), 47–57.

[25] Thomson, J. D. Effects of variation in inflorescence size and floral rewards on the visitation rates of traplining pollinators of Aralia hispida. Evol. Ecol. 2 (1988), 65–76.

[26] Andersson, S. Floral display and pollination success in Senecio jacobaea (Asteraceae): interactive effects of head and corymb size. Am. J. Bot. 83 (1996), 71–75

[27] Kirchner, F., S. H. Luijten, E. Imbert, M. Riba, M. Mayol, S. C. Gonza, and B. Colas. Effects of local density on insect visitation and fertilization success in the narrow-endemic Centaurea corymbosa (Asteraceae). Oikos 111 (2005), 130–142.

[28] Iwata, T., O. Nagasaki, H. S. Ishii, and A. Ushimaru. Inflorescence architecture affects pollinator behaviour and mating success in Spiranthes sinensis (Orchidaceae). New Phytol. 193 (2012), 196–203.

[29] Fenner, M., J. E. Cresswell, R. A. Hurley, and T. Baldwin. Relationship between capitulum size and pre-dispersal seed predation by insect larvae in common Asteraceae. Oecologia 130 (2002), 72–77.

[30] Fenner, M., W. G. Lee, and E. H. Pinn. Reproductive features of Celmisia species (Asteraceae) in relation to altitude and geographical range in New Zealand. Biol. J. Linn. Soc. 74 (2001), 51–58.

[31] Sargent, R. D., C. Goodwillie, S. Kalisz, and R. H. Ree. Phylogenetic evidence for a flower size and number trade-off. Am. J. Bot. 94 (2007), 2059–2062.

[32] Goodwillie, C., R. D. Sargent, C. G. Eckert, E. Elle, M. A. Geber, M. O. Johnston, S. Kalisz, D. A. Moeller, R. H. Ree, M. Vallejo-Marín, and A. A. Winn. Correlated evolution of mating system and floral display traits in flowering plants and its implications for the distribution of mating system variation. New Phytol. 185 (2010), 311–321.

[33] Vasconcelos, T. N. C., and C. E. B. Proença. Floral cost vs. floral display: Insights from the megadiverse Myrtales suggest that energetically expensive floral parts are less phylogenetically constrained. Am. J. Bot. 102 (2015), 900–909.

[34] Funk, V. A., A. Susanna, T. Stuessy, and R. J. Bayer, eds. Systematics, evolution, and biogeography of Compositae. International Association for Plant Taxonomy, Viena, 2009.

[35] Harris, E. M. Inflorescence and floral ontogeny in Asteraceae: a synthesis of historical and current concepts. Bot. Rev. 61 (1995), 93–278.

[36] Pozner, R., C. Zanotti, and L. A. Johnson. Evolutionary origin of the Asteraceae capitulum: Insights from Calyceraceae. Am. J. Bot. 99 (2012), 1–13.

[37] Torices, R., M. Méndez, and J. M. Gómez. Where do monomorphic sexual systems fit in the evolution of dioecy? Insights from the largest family of angiosperms. New Phytol. 190 (2011), 234–48.

[38] Alkio, M., and E. Grimm. Vascular connections between the receptacle and empty achenes in sunflower (Helianthus annuus L.). J. Exp. Bot. 54 (2003), 345–348.

[39] Alkio, M., A. Schubert, W. Diepenbrock, and E. Grimm. Effect of source – sink ratio on seed set and filling in sunflower (Helianthus annuus L.). Plant. Cell Environ. 26 (2003), 1609–1619.

[40] Stephenson, A. G. The regulation of maternal investment in plants. Pp. 151–171 in C. Marshall, J. Grace eds. Fruit and seed production: aspects of development, environmental physiology and ecology. Cambridge University Press, Cambridge, 1992.

[41] Funk, V. A., L. Watson, B. Gemeinholzer, E. Schilling, A. Susanna, and R. K. Jansen. Everywhere but Antarctica: using a supertree to understand the diversity and distribution of the Compositae. Biol. Skr. 55 (2005), 343–374.

[42] Schneider, C. A., W. S. Rasband, and K. W. Eliceiri. NIH Image to ImageJ: 25 years of image analysis. Nat. Methods 9 (2012), 671–5.

[43] Harmon, L. J., and J. B. Losos. The effect of intraspecific sample size on type I and type II error rates in comparative studies. Evolution 59 (2005), 2705–2710.

[44] Freckleton, R. P., P. H. Harvey, and M. Pagel. Phylogenetic analysis and comparative data: a test and review of evidence. Am. Nat. 160 (2002), 712–26.

[45] Paradis, E. Analysis of phylogenetics and evolution with R (Second Edition). Springer. New York, 2012.

[46] Felsenstein, J. Inferring phylogenies. Sinauer Associates, Sunderland, 2004.

[47] Paradis, E., J. Claude, and K. Strimmer. APE: Analyses of phylogenetics and evolution in R language. Bioinformatics 20 (2004), 289–290.

[48] Harmon L. J, J. T. Weir, C. D. Brock, R. E. Glor, and W. Challenger. GEIGER: investigating evolutionary radiations. Bioinformatics 24 (2008), 129–131.

[49] Torices, R. Adding time-calibrated branch lengths to the Asteraceae supertree. J. Syst. Evol. 48 (2010), 271–278.

[50] Lenth, R. V. Least-squares means: the R package lsmeans. J Stat Soft 69 (2016), 1–33.

[51] R Core Team. R: A language and environment for statistical computing. R Foundation for Statistical Computing, Vienna, Austria, 2015. http://www.R-project.org/

[52] Schwarzer, G. meta: general package for meta-analysis. R package version 4.1–0, 2015. http://CRAN.R-project.org/package=meta

[53] Revell, L. J. phytools: An R package for phylogenetic comparative biology (and other things). Methods Ecol. Evol. 3 (2012), 217–223.

[54] Harder, L. D., S. C. H. Barrett, and W. W. Cole. The mating consequences of sexual segregation within inflorescences of flowering plants. Proc. R. Soc. B Biol. Sci. 267 (2000), 315–20.

[55] Harder, L. D., and S. C. H. Barrett. Mating cost of large floral displays in hermaphrodite plants. Nature 373 (1995), 512–515.

[56] Marshall, D. F., and R. J. Abbott. Polymorphism for outcrossing frequency at ray floret locus in Senecio vulgaris L. III. Causes. Heredity 53 (1984), 145–149.

[57] Cheptou, P. O., J. Lepart, and J. Escarre. Differential outcrossing rates in dispersing and non-dispersing achenes in the heterocarpic plant Crepis sancta (Asteraceae). Evol. Ecol. 15 (2001), 1–13.

[58] Gibson, J. Ecological and genetic comparison between ray and disc achene pools of the heteromorphic species Prionopsis ciliata (Asteraceae). Int. J. Plant Sci. 162 (2001), 137–145.

[59] Gibson, J. P., A. D. Tomlinson, and A. S. College. Genetic diversity and mating system comparisons between ray and disc achene seed pools of the heterocarpic species Heterotheca subaxillaris (Asteraceae). Int. J. Plant Sci. 163 (2002), 1025–1034.

[60] Ferrer, M. M., and S. V Good-Avila. Macrophylogenetic analyses of the gain and loss of self-incompatibility in the Asteraceae. New Phytol. 173 (2007), 401–14.

[61] Schoen, D. J., and M. Dubuc. The evolution of inflorescence size and number: a gamete-packaging strategy in plants. Am. Nat. 135 (1990), 841–857.

[62] Fishbein, M., and D. L. Venable. Evolution of inflorescence design: theory and data. Evolution 50 (1996), 2165–2177.

[63] Torices, R., and A. A. Anderberg. Phylogenetic analysis of sexual systems in Inuleae (Asteraceae). Am. J. Bot. 96 (2009), 1011–1019.

[64] Sutherland, S. Patterns of fruit-set: what controls fruit-flower ratios in plants? Evolution 40 (1986), 117–128.

[65] Vamosi, S. M., S. J. Mazer, and F. Cornejo. Breeding systems and seed size in a neotropical flora: testing evolutionary hypotheses. Ecology 89 (2008), 2461–72.

[66] Reznick, D., L. Nunney, and A. Tessier. Big houses, big cars, superfleas and the costs of reproduction. Trends Ecol. Evol. 15 (2000), 421–425.

[67] Herrera, C. M. Multiplicity in unity: plant subindividual variation and interactions with animals. The University of Chicago Press, London, UK, 2009.

