## Supplementary Figure S1 for "Architectural traits constrain the evolution of unisexual flowers and sexual segregation within inflorescences: an interspecific approach"

**Supplementary Figure S1. Phylogenetic distribution of sampled species.** The number of species sampled in each lineage are displayed on the right of the phylogenetic tree showing major lineages of Asteraceae (based on Funk's et al. (2005) supertree). The color of the number indicates the sexual system of the species: grey, hermaphroditic; black, gynomonoecious; and, red, monoecious. Branch color also denotes the most likely ancestral sexual system for each lineage (based on Torices et al. (2011)).

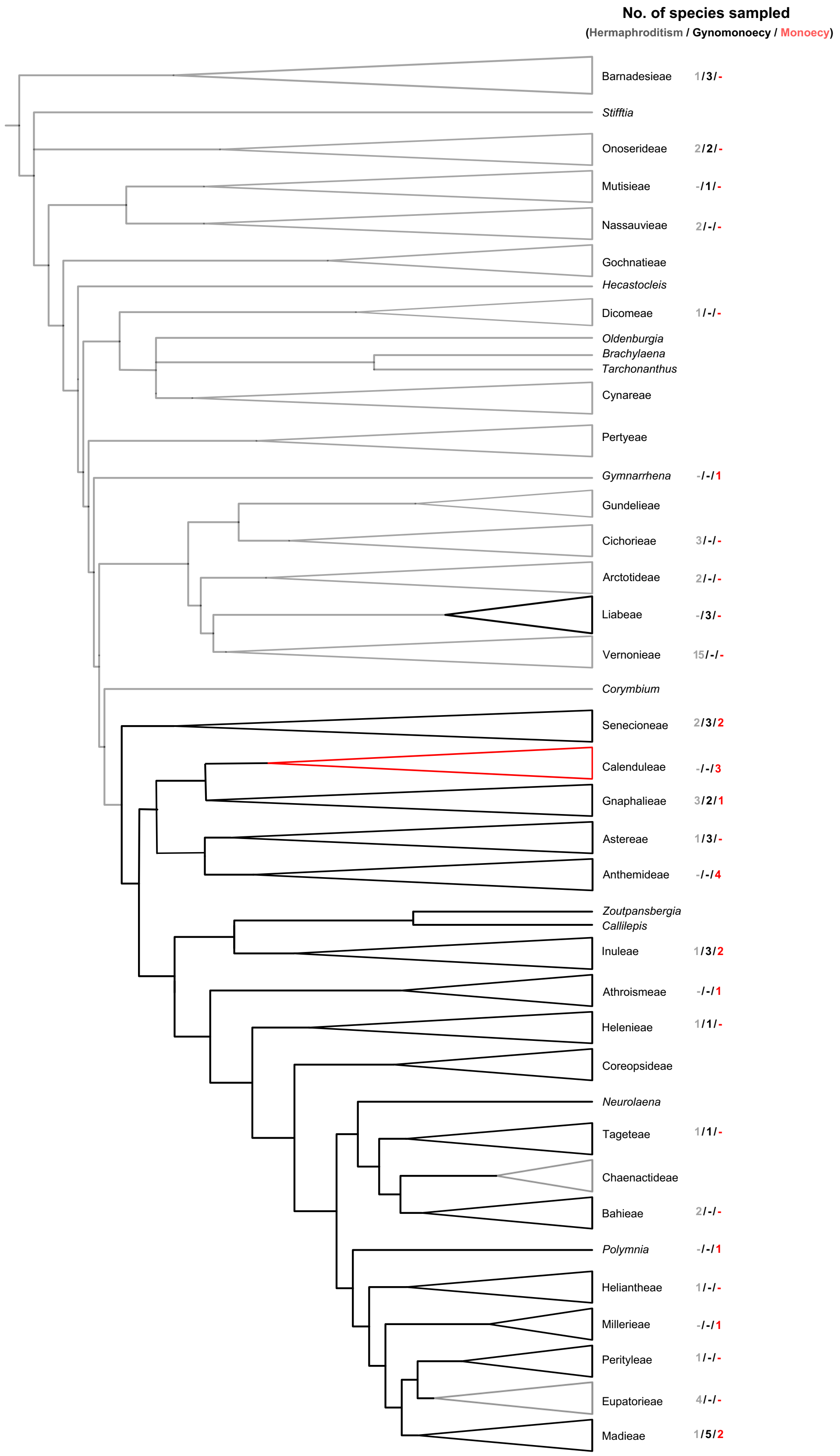
