## Supplementary Tables S1-5 for "Architectural traits constrain the evolution of unisexual flowers and sexual segregation within inflorescences: an interspecific approach"

**Supplementary Table S1**. Average ± SD size (estimated as area in $mm^2$) of outer and inner fruits of gynomonoecious, monoecious and hermaphrotidic Asteraceae species. FSD – Fruit size difference measured as meta-analytical effect size. [1]Herbarium: Swedish Museum of Natural History (S); Herbarium of University of Coimbra (COI). [2]Dist.: major areas where the species is distributed. [3]Sexual system: H – Hermaphroditism; GM – Gynomonoecy; M – Monoecy.

| Subfamily | Tribe | Species | Herbarium[1] | Herbarium Voucher | Dist.[2] | Sexual System[3] | Outermost fruits | | Innermost fruits | | |
|---|---|---|---|---|---|---|---|---|---|---|---|
| | | | | | | | n | Mean ± SD | n | Mean ± SD | FSD ± SE |
| Asteroideae | Anthemideae | *Artemisia crithmifolia* | COI | Aarão F. de Lacerda, no. 751-780 | Europe | M | - | - | - | - | - |
| | | *Eriocephalus umbellatus* | COI | Heron, s.n. | South Africa | M | 1 | 1.82 | - | - | - |
| | | *Hippia fruticosa* | COI | Lason, no. 10686 | South Africa | M | 10 | 1.63±0.41 | | | |
| | | *Soliva pterosperma* | COI | J. Matos; A Matos & A. Marques, no. 750-4806 | South America | M | 10 | 1.55±0.25 | 5 | 1.02±0.21 | 2.092±0.706 |
| | Astereae | *Amellus strigosus* | S | E. Wall, no. 137 | South Africa | GM | 33 | 2.27±0.16 | 15 | 2.14±0.20 | 0.738±0.321 |
| | | *Calotis erinaceae* | S | E.N.S. Jackson, no. 5948 | Australia | GM | 38 | 3.24±0.51 | 12 | 3.14±0.43 | 0.200±0.332 |
| | | *Grindelia arenicola* | S | E. K. Balls, no. 10161 | North America | GM | 54 | 3.36±0.45 | 25 | 3.86±0.56 | -1.017±0.256 |
| | | *Pteronia incana* | S | A. & B. Strid, no. 37701 | South Africa | H | 8 | 4.41±0.65 | - | - | - |
| | Athroismeae | *Blepharispermum spinulosum* | COI | Cyossmeiler, no. 8059 | Central Africa | M | 1 | 4.64 | - | - | - |
| | Calenduleae | *Calendula arvensis* | COI | J. Nogueirs, no. 757-10962 | Europe, Mediterranean Basin | M | 6 | 17.47±5.02 | 8 | 8.98±0.47 | 2.438±0.766 |

| Subfamily | Tribe | Species | Herbarium[1] | Herbarium Voucher | Dist.[2] | Sexual System[3] | Outermost fruits | | Innermost fruits | | FSD ± SE |
|---|---|---|---|---|---|---|---|---|---|---|---|
| | | | | | | | n | Mean ± SD | n | Mean ± SD | |
| | | *Dimorphotheca sinuata* | COI | Sange Kloof, no. 8598 | South Africa | M | - | - | - | - | - |
| | | *Osteospermum hispidum* | COI | Elands, no. 9755 | South Africa | M | - | - | - | - | - |
| | Gnaphalieae | *Anaxeton arborescens* | S | A. Meelbold, no.13494 | South Africa | M | - | - | - | - | - |
| | | *Ammobium alatum* | S | A. Anderberg & A.L. Anderberg, no. 7148 | Australia | H | 53 | 1.25±0.24 | 88 | 1.08±0.27 | 0.652±0.178 |
| | | *Millotia myosotidifolia* | S | F.J. Badman, no. 8397 | Australia | H | 44 | 0.84±0.18 | 21 | 1.07±0.23 | -1.152±0.285 |
| | | *Ozothamnus diosmifolius* | S | A Anderberg & A.-L Anderberg, no. 7043 | Australia | H | - | - | - | - | - |
| | | *Plecostachys serpyllifolia* | S | R. D. A. Bayliss, no. 8375 | South Africa | GM | 10 | 1.56±0.09 | - | - | - |
| | | *Rosenia humilis* | S | Kare Bremer, no. 164 | South Africa | GM | 10 | 2.45±0.32 | 8 | 2.23±0.23 | 0.737±0.494 |
| | Inuleae | *Blumea riparia* | S | Chieng-Chang Hsu, no. 5201 | Eastern Asia | GM | 172 | 0.39±0.05 | 44 | 0.34±0.05 | 0.207±0.166 |
| | | *Epaltes cunninghamii* | S | B. Nordenstam & A. Anderberg, no. 972 | Australia | M | 16 | 0.43±0.19 | 11 | 0.42±0.08 | 0.062±0.392 |
| | | *Inula oculus-christi* | S | I. Segelberg, no. 13761/5 | Europe | GM | 26 | 0.88±0.14 | 75 | 0.97±0.11 | -0.755±0.234 |
| | | *Inula peacockiana* | S | K. H. Rechinger, no. 49051 | Western Asia | H | 16 | 5.63±0.49 | 4 | 5.52±0.43 | 0.219±0.560 |
| | | *Pluchea dentex* | S | B. Nordenstam & A. Anderberg, no. 325 | Australia | M | 57 | 0.18±0.04 | 81 | 0.18±0.03 | 0.000±0.173 |
| | | *Streptoglossa liatroides* | S | A. Strid, no. 4269 | Australia | GM | 34 | 1.79±0.21 | 27 | 1.86±0.18 | -0.350±0.260 |

| Subfamily | Tribe | Species | Herbarium[1] | Herbarium Voucher | Dist.[2] | Sexual System[3] | Outermost fruits | | Innermost fruits | | FSD ± SE |
|---|---|---|---|---|---|---|---|---|---|---|---|
| | | | | | | | n | Mean ± SD | n | Mean ± SD | |
| | Senecioneae | *Blennosperma californicum* | | Lewis S. Rose, no. 9308-33008 | North America | M | 3 | 2.25±0.11 | - | - | - |
| | | *Kleinia longiflora* | S | E. Wall, no. 622 | Africa | H | 8 | 6.74±1.19 | 3 | 7.46±0.28 | -0.622±0.697 |
| | | *Ligularia fischeri* | S | M Mizushim, no. 13766 | Eastern Asia | GM | 11 | 4.96±0.50 | 7 | 4.97±0.65 | -0.017±0.484 |
| | | *Othonna coronopifolia* | | Iaron, no. 7885 | South Africa | M | 7 | 1.85±0.32 | - | - | - |
| | | *Roldana mexicana* | S | Geo. B. Hinton, no 8745 | Mexico | H | 14 | 1.42±0.24 | 5 | 1.37±0.26 | 0.195±0.522 |
| | | *Senecio inornatus* | S | DM Hilliard & B.L. Burtt, no. 7492 | Africa | GM | 22 | 0.66±0.14 | 7 | 0.74±0.05 | -0.619±0.443 |
| | | *Senecio subsessilis* | S | J.A. Mlangwa , P.B. Phillipson, H. van Vlaenderen & W. Kindeketa, no. 305 | Africa | GM | 20 | 3.16±0.35 | 8 | 3.38±0.45 | -0.563±0.426 |
| Barnadesioideae | Barnadesieae | *Barnadesia spinosa* | S | H. Humbert, no. 26923 | South America | H | 12 | 3.89±0.59 | 3 | 5.17±1.78 | -1.362±0.708 |
| | | *Dasyphyllum diacanthoides* | S | Mleyer, no. 8161 | South America | GM | 14 | 2.56±0.38 | 4 | 2.25±0.58 | 0.695±0.582 |
| | | *Dasyphyllum ferox* | S | C. Hammarlund, no. 534 | South America | GM | 7 | 4.10±0.38 | 5 | 4.35±0.49 | -0.540±0.601 |
| | | *Doniophyton anomalon* | S | F. Barkley & O. Paci, s.n. | South America | GM | 31 | 12.41±1.63 | 15 | 13.99±1.47 | -0.982±0.332 |
| Carduoideae | Dicomeae | *Dicoma anomala* | S | H. & HE. Wanntorp, no. 464 | Africa | H | 9 | 2.37±0.33 | 4 | 2.75±0.43 | -0.982±0.644 |
| Cichorioideae | Arctotideae | *Hirpicium echinus* | S | Lars Erik Kers, no. 2179 | South Africa | H | 11 | 2.28±0.44 | 5 | 2.24±0.39 | 0.089±0.540 |
| | | *Hoplophyllum spinosoum* | S | P. Goldblatt, no. 4325 | South Africa | H | 7 | 4.69±0.76 | | | |

| Subfamily | Tribe | Species | Herbarium[1] | Herbarium Voucher | Dist.[2] | Sexual System[3] | Outermost fruits | | Innermost fruits | | FSD ± SE |
|---|---|---|---|---|---|---|---|---|---|---|---|
| | | | | | | | n | Mean ± SD | n | Mean ± SD | |
| | Cichorieae | *Microseris douglasii* | S | Lewis S. Rose, no. 66037B | North America | H | 25 | 2.33±0.14 | 15 | 2.19±0.19 | 0.856±0.342 |
| | | *Uropappus lindleyi* | S | L.S. Rose, no. 63059 | North America | H | 23 | 4.54±0.13 | 9 | 4.81±0.13 | -2.025±0.477 |
| | | *Warionia saharae* | S | E.K. Balls, no. 2530 | North Africa | H | 21 | 15.31±2.69 | 19 | 15.51±2.49 | -0.075±0.317 |
| | Liabeae | *Liabum bourgeaui* | S | Robert Merrill King & Victor Castro, no. 9997 | Mesoamerica | GM | 52 | 0.15±0.03 | 17 | 0.17±0.03 | -0.659±0.285 |
| | | *Philoglossa peruviana* | S | E. Asplund, no. 13735 | South America | GM | 15 | 1.10±0.08 | 4 | 1.08±0.07 | 0.244±0.564 |
| | | *Sinclairia polyantha* | S | C. L. Lundell & Elias Contreras, no 20619 | Mesoamerica | GM | 14 | 0.43±0.07 | 4 | 0.49±0.11 | -0.723±0.583 |
| | Vernonieae | *Baccharoides adoensis* | S | M Reekmans, no. 9172 | Africa | H | 29 | 4.47±0.55 | 18 | 4.57±0.59 | -0.174±0.301 |
| | | *Centrapalus pauciflorus* | S | T. Eriksson, V. Kalema & G. Leliyo, no. TE 533 | Africa | H | 22 | 3.62±0.28 | 14 | 3.73±0.42 | -0.316±0.344 |
| | | *Critoniopsis leiocarpa* | S | Ynes Mexia, No. 9119 | Mesoamerica | H | - | - | - | - | - |
| | | *Cyanthillium cinereum* | S | Dick Hummel, s.n | Tropical Asia | H | 13 | 0.70±0.05 | 6 | 0.76±0.17 | -0.566±0.504 |
| | | *Ethulia conyzoides* | S | H.J. Venter & A. Venter, no. 9677 | Africa | H | 15 | 0.88±0.14 | 7 | 0.85±0.03 | 0.244±0.460 |
| | | *Gymnanthemum amygdalinum* | S | Fernandez Casas, no. 11433 | Africa | H | 7 | 1.74±0.29 | 5 | 1.57±0.19 | 0.616±0.605 |
| | | *Lepidaploa tortuosa* | S | Llewelyn Willians, s.n. | Meso America | H | 22 | 1.40±0.32 | 11 | 1.12±0.31 | 0.862±0.386 |
| | | *Linzia glabra* | S | E. Lawrence, no. 112 | Africa | H | 8 | 4.58±0.61 | 2 | 4.36±0.19 | 0.346±0.797 |
| | | *Orbivestus cinerascens* | S | Lars Erik Kers, no. 593 | Africa | H | 10 | 1.22±0.14 | 5 | 1.08±0.22 | 0.781±0.572 |

| Subfamily | Tribe | Species | Herbarium[1] | Herbarium Voucher | Dist.[2] | Sexual System[3] | Outermost fruits | | Innermost fruits | | FSD ± SE |
|---|---|---|---|---|---|---|---|---|---|---|---|
| | | | | | | | n | Mean ± SD | n | Mean ± SD | |
| | | *Parapolydora fastigiata* | S | O.H. Volk, no. 00367 | Africa | H | 36 | 2.26±0.55 | 19 | 1.66±0.31 | 1.227±0.308 |
| | | *Polydora poskeana* | S | E.S. Pooley, no. 477 | Africa | H | 30 | 1.90±0.22 | 13 | 1.91±0.36 | -0.037±0.332 |
| | | *Vernonanthura patens* | S | E. Wall, no. 9301 | Central and South America | H | 9 | 0.90±0.09 | 3 | 0.82±0.14 | 0.724±0.691 |
| | | *Vernonanthura alamanii* | S | H. Fröderström & E. Hultén, no. 321 | Mesoamerica | H | 46 | 4.10±0.69 | 25 | 4.43±0.76 | -0.456±0.252 |
| | | *Vernonia angustifolia* | S | Ted Bradley, no. 3502 | North America | H | 13 | 1.69±0.31 | 5 | 1.69±0.23 | 0.000±0.526 |
| | | *Vernonia lasiopus* | S | T. Erikson, V. Kalerna & G. Leliyo, no. TE 546 | Africa | H | 10 | 1.38±0.28 | 5 | 0.93±0.24 | 1.578±0.642 |
| Gymnarrhenoideae | Gymnarrheneae | *Gymnarrhena micrantha* | COI | A. Grizi, no. 8970-383 | Middle East | M | 13 | 1.31±0.40 | 7 | 1.60±0.24 | -0.783±0.489 |
| Mutisioideae | Mutisieae | *Chaptalia nutans* | S | E. Wall, no. 729 | North and South America | GM | 32 | 2.57±0.37 | 19 | 2.90±0.26 | -0.973±0.307 |
| | Nassauvieae | *Jungia paniculata* | S | S.G. Saunders, no. 1244 | South America | H | 19 | 0.69±0.12 | 8 | 0.62±0.12 | 0.566±0.430 |
| | | *Perezia multiflora* | S | Kjell von Sneiden, no. A333 | South America | H | 22 | 3.78±0.48 | 24 | 3.73±0.61 | 0.089±0.295 |
| | Onoserideae | *Onoseris alata* | S | J. Olea, s.n. | South America | GM | 16 | 4.42±0.90 | 9 | 3.14±0.46 | 1.596±0.484 |
| | | *Onoseris odorata* | S | Francis W Pennell, no 14468 | South America | GM | 20 | 3.41±0.35 | 10 | 3.78±0.50 | -0.890±0.406 |
| | | *Plazia argentea* | S | E. Carrette, s.n. | South America | H | 4 | 4.55±1.17 | 1 | 4.42 | - |

| Subfamily | Tribe | Species | Herbarium[1] | Herbarium Voucher | Dist.[2] | Sexual System[3] | Outermost fruits | | Innermost fruits | | |
|---|---|---|---|---|---|---|---|---|---|---|---|
| | | | | | | | n | Mean ± SD | n | Mean ± SD | FSD ± SE |
| | | *Trixis antimenorrhoea* | S | F.J. Breteler, no. 3502 | South America | H | 7 | 1.23±0.04 | 2 | 1.21±0.14 | 0.275±0.806 |
| "Heliantheae alliance" | Bahieae | *Florestina pedata* | S | Maury, no.24 | Mesoamerica | H | 6 | 2.25±0.28 | 4 | 1.85±0.19 | 1.445±0.767 |
| | | *Palafoxia arida* | S | K. Bremer, no. 2479 | North America | H | 12 | 7.95±1.31 | 5 | 7.64±1.01 | 0.238±0.534 |
| | Eupatorieae | *Ageratina calaminthaefolia* | S | Robert Merrill King & Paul M. Peterson, no. 9957 | North America | H | 7 | 1.14±0.18 | 3 | 1.33±0.31 | -0.781±0.726 |
| | | *Brickellia chlorolepis* | S | Robert Merrill King & Paul M. Peterson, no.9836 | North America | H | 13 | 2.10±0.26 | 9 | 1.88±0.45 | 0.607±0.445 |
| | | *Chromolaena odorata* | S | Erik Wall, no.72 | North America | H | 17 | 1.36±0.18 | 13 | 1.40±0.21 | -0.201±0.369 |
| | | *Liatris aspera* | S | D.S. Correll & H. B. Correll, no. 36587 | North America | H | 17 | 4.15±0.37 | 8 | 4.49±0.36 | -0.896±0.450 |
| | Helenieae | *Baileya pleniradiata* | S | J. Laubert, no 113 | North America | GM | 100 | 1.49±0.18 | 25 | 1.37±0.14 | 0.690±0.228 |
| | | *Marshallia graminifolia* | S | C. Ritchie Bell, no. 15744 | North America | H | 26 | 3.12±0.32 | 29 | 3.61±0.35 | -1.437±0.305 |
| | Heliantheae | *Rudbeckia fulgida* | S | F.T. McFarland, no. 347 | North America | H | 51 | 1.15±0.14 | 32 | 1.16±0.10 | -0.079±0.226 |
| | Madieae | *Anisocarpus scabridus* | S | M.S. Baker, no 10658 | North America | GM | 4 | 11.02±0.70 | 1 | 8.81 | - |
| | | *Arnica lanceolata* | S | Galen Smith, no. 2049 | North America | GM | 43 | 3.20±0.49 | 7 | 4.51±0.48 | -2.638±0.492 |
| | | *Dubautia laxa* | S | L.M. Cranwell, no. 3417 | Hawaii | H | 8 | 1.24±0.21 | 2 | 1.79±0.10 | -2.489±1.066 |
| | | *Hemizonia fasciculata* | COI | S.B. & W.F. Parish, no. 9254 | North America | M | 5 | 1.67±0.08 | - | - | - |

| Subfamily | Tribe | Species | Herbarium[1] | Herbarium Voucher | Dist.[2] | Sexual System[3] | Outermost fruits | | Innermost fruits | | FSD ± SE |
|---|---|---|---|---|---|---|---|---|---|---|---|
| | | | | | | | n | Mean ± SD | n | Mean ± SD | |
| | | *Holozonia filipes* | COI | S.B. & W.F. Parish, no. 9257-486 | North America | M | - | - | - | - | - |
| | | *Layia platyglossa* | COI | William H. Beble, no. 9258 | North America | GM | 20 | 1.93±0.17 | 15 | 1.98±0.17 | -0.287±0.344 |
| | | *Madia anomala* | S | David. D. Keck, no. 2313 | North America | GM | - | - | - | - | - |
| | | *Monolopia lanceolata* | S | E.K. Balls, no. 8547 | North America | GM | 22 | 1.05±0.15 | 10 | 0.99±0.17 | 0.374±0.385 |
| | Millerieae | *Melampodium leucanthum* | COI | W.P. Cottam, no. 9129-10231 | North America | M | 7 | 2.58±0.74 | - | - | - |
| | Perityleae | *Perityle emoryi* | S | M.O. Dillon & D.O. Dillon, no. 4850 | North and South America | GM | 44 | 1.63±0.14 | 23 | 1.59±0.24 | 0.219±0.258 |
| | Polymnieae | *Polymnia canadensis* | | Grady L. & Barbara D. Webter, no. 9122-7088 | North America | M | 4 | 5.12±0.40 | - | - | - |
| | Tageteae | *Oxypappus scaber* | S | Mexia, no. 1367 | Mesoameri ca | GM | 22 | 0.17±0.03 | 8 | 0.16±0.02 | 0.350±0.416 |
| | | *Porophyllum scoparium* | S | K. Bremer, no. 2379 | North America | H | 39 | 1.56±0.23 | 24 | 1.82±0.21 | -1.153±0.280 |

**Supplementary Table S2.** Phylogenetic regression between inflorescence size (head diameter), number of flowers and flower density (number of flowers / $mm^2$). All variables were log transformed.

| | Inflorescence size | | | No. of flowers | | |
|---|---|---|---|---|---|---|
| | *b* ± SE | *t* | *P* | *b* ± SE | *t* | *P* |
| No. of flowers | 0.72 ± 0.13 | 5.73 | 0.000 | - | - | - |
| Flower density | -1.25 ± 0.13 | -0.74 | 0.000 | 0.24 ± 0.13 | 1.81 | 0.073 |

**Supplementary Table S3.** Phylogenetic RMA regressions between inflorescence size (head diameter), number of flowers and flower density (number of flowers / $mm^2$). All variables were log transformed. Estimated slopes were tested against the null hypothesis of $b = 1$.

| | Inflorescence size | | | | No. of flowers | | | |
|---|---|---|---|---|---|---|---|---|
| | *b* | *t* | *d.f.* | *P* | *b* | *t* | *d.f.* | *P* |
| No. of flowers | 1.42 | 3.94 | 74.9 | 0.0002 | - | - | - | - |
| Flower density | -1.65 | 6.64 | 69.7 | <0.0001 | 1.17 | 1.44 | 85.9 | 0.153 |

**Supplementary Table S4.** Phylogenetic regression between FSD, outer and inner fruit size and inflorescence size (head diameter), number of flowers and flower density (number of flowers / $mm^2$). All variables were log transformed.

| | FSD | | | Outer fruit size | | | Inner fruit size | | |
|---|---|---|---|---|---|---|---|---|---|
| **Inflorescence traits** | *b* ± SE | *t* | *P* | *b* ± SE | *t* | *P* | *b* ± SE | *t* | *P* |
| Head diameter (mm) | -1.15 ± 0.40 | -3.11 | 0.003 | 0.69 ± 0.13 | 5.43 | < 0.001 | 0.89 ± 0.14 | 6.54 | < 0.001 |
| No. of flowers | -0.14 ± 0.28 | -0.50 | 0.621 | - 0.16 ± 0.10 | -1.53 | 0.130 | -0.14 ± 0.12 | -1.12 | 0.267 |
| Flower density (no. flowers/$mm^2$) | 0.46 ± 0.19 | 2.45 | 0.017 | - 0.60 ± 0.05 | -10.57 | < 0.001 | -0.66 ± 0.06 | -11.23 | < 0.001 |

**Supplementary Table S5.** Deviance analysis of the phylogenetic generalized linear model fitting Fruit Size Difference (FSD) as the response of the degree of floral sexual specialization within inflorescence (hermaphroditism, gynomonoecy or monoecy), and flower density. FSD is the standardized fruit size difference between outer and inner fruits measured as the meta-analytical effect size. Flower density was log-transformed.

| **Variables** | *F* | d.f. | *P* |
|---|---|---|---|
| Intercept | 0.77 | 1, 62 | 0.385 |
| Degree of floral sexual specialization | 0.87 | 2, 62 | 0.423 |
| Flower density (no. flowers/mm$^2$) | 4.36 | 1, 62 | 0.041 |
